# Integration of an anellovirus genome in the SKNO-1 acute myeloid leukemia cell line

**DOI:** 10.64898/2026.01.22.701047

**Authors:** Na Cui, Stephanie Goya, Eli A. Piliper, Alexander L. Greninger

**Author notes:** Corresponding author Alexander L. Greninger, Department of Laboratory Medicine and Pathology, University of Washington Medical Center 850 Republican St, Seattle, WA 98109, USA.

## Abstract

Anelloviruses are highly diverse, ubiquitous single-stranded DNA viruses whose replication, cellular reservoirs, and mechanisms of persistence remain poorly understood. Here, we identify and characterize a Alphatorquevirus homin 29 genome stably integrated into an rDNA locus of human chromosome 21 in the acute myeloid leukemia cell line SKNO-1. Large-scale mining of NCBI Sequence Read Archive datasets revealed unusually high anellovirus k-mer abundance specifically in sequencing runs of the SKNO-1 cell line from multiple laboratories. De novo assembly of RNA-seq data recovered a 3.25 kb viral genome, and long-read PacBio sequencing confirmed its integration within the *RNA45SN2* gene on human chromosome 21. Digital droplet PCR (ddPCR) quantified ∼0.5 viral genomes per cell, and RT-ddPCR detected transcripts from ORF1 and viral terminal repeats, consistent with transcription of the integrant. The ORF1 coding sequence encoded for a truncated but structurally conserved capsid protein retaining jelly roll and P-domain features but lacking the C-terminal domain. Analysis of public ChIP-seq data from the SKNO-1 cell line demonstrated broad occupancy of the BRD4 transcriptional coactivator across the viral genome along with focal enrichment of hematopoietic ETS family transcription factors within the integrant’s ∼300 bp untranslated region upstream of viral open reading frames. Together, these findings demonstrate stable integration, chromatin accommodation, and transcriptional maintenance of an Alphatorquevirus in a human leukemia cell line, providing a unique model to study host tolerance and transcriptional regulation of anellovirus DNA and informing our understanding of anellovirus hematopoietic cell tropism.

**IMPORTANCE:** Anelloviruses are a curious group of highly genetically diverse single-stranded DNA viruses that have been found ubiquitously among humans and are hypothesized to be a potential commensal human virus. Their omnipresence has been matched only by the dearth of our understanding of their basic biological processes – how they persist, how they replicate, and how they spread. Here we identify a naturally integrated Alphatorquevirus genome in the widely used AML cell line SKNO-1 and show that it is stably maintained, transcribed, and embedded within a rDNA locus on human chromosome 21. This discovery highlights the ability to anelloviruses to integrate into human DNA, generates hypotheses around transcription factors used in anellovirus gene transcription, and suggests anelloviruses can persist in myeloblasts.

## INTRODUCTION

Anelloviruses comprise a highly diverse group of small, circular, single-stranded DNA viruses that are ubiquitous in humans yet poorly understood in terms of their biology and clinical significance. Since their discovery in 1997, anelloviruses have been detected in approximately 90% of adults and are most commonly found in blood and bone marrow (1). Viral DNA has also been identified in multiple human tissues and body fluids, including ovary, lung, saliva, and respiratory secretions (2, 3).

Advances in viral metagenomics have expanded the known diversity of human anelloviruses and renewed interest in their diagnostic and therapeutic potential (3). Studies of transfusion- and transplantation-associated infections have demonstrated that donor-derived anellovirus DNA can persist in recipients for several months (4, 5). Moreover, circulating anellovirus have been proposed as potential biomarkers for organ transplant rejection (6–9). Despite these associations, the mechanism underlying anellovirus persistence, replication, and immune modulation remain largely unclear (10–12).

Anelloviruses exhibit substantial genetic diversity approaching that of HIV-1 (4). As of 2025, the International Committee on Taxonomy of Viruses (ICTV) classifies the *Anelloviridae* family into 34 genera and 173 species identified across diverse mammal host (13–17). Human anelloviruses are currently assigned to three genera: *Alphatorquevirus* (Torque Teno viruses, TTV), *Betatorquevirus* (Torque Teno mini viruses, TTMV), and *Gammatorquevirus* (Torque Teno midi viruses, TTMDV) (3). Members of this family are non-enveloped and contain a 2.0-3.9 kb negative-sense circular DNA genome encoding six open reading frames (ORF) through alternative splicing (18–20). Because they lack a viral DNA polymerase, anelloviruses depend on host replication machinery, likely replicating in the cell nucleus through a rolling-circle mechanism (10, 21). Although human anelloviruses have been detected in a variety of tissues, the primary cellular reservoir supporting active replication remains uncertain. Evidence from previous studies suggests replication may occur in bone marrow cells and activated peripheral blood mononuclear cells (22, 23). However, progress in defining viral tropism and replication dynamics has been constrained by the absence of efficient cell culture systems or suitable animal models. To date, most insights have derived from plasmid-based transfection studied in human cell lines such as COS1, HEK293, and L428, which do not support productive viral replication (19, 24).

In this study, we describe the discovery of an Alphatorquevirus in the human acute myeloid leukemia (AML) cell line SKNO-1. Remarkably, the viral genome was integrated into the host *RNA45SN2* gene on chromosome 21 and stably transcribed and maintained in the cell line. Through combined bioinformatic and experimental approaches, we confirm the presence and transcriptional activity of this anellovirus and use existing systems biology datasets using this cell line to inform host cell tropism and anellovirus transcription.

## RESULT

### Anellovirus k-mers are primarily detected in public sequencing data from SKNO-1 cell line

We investigated the presence of anellovirus in public sequencing data by querying the Cloud-based Taxonomy Analysis Table on NCBI SRA using BigQuery. This search yielded 218,015 SRA datasets with at least two anellovirus k-mer hits of 32 nt in length (Table S1). To identify a potential model cell line for anellovirus studies, we cross-referenced the SRA accessions with a dataset of the 333,979 SRA datasets annotated as a human cell line (HCL) (Table S2). We identified 5,438 overlapping HCL datasets that contained at least two anellovirus k-mer hits (Table S3, Figure 1A). Of these, the vast majority (4,959/5,438) had 2-9 anellovirus k-mer hits, while 385 HCL runs contained 10-100 hits, and notably, 94 HCL runs – 1.7% of anellovirus-containing HCL datasets – had >100 k-mer hits, contributing 83.9% of the total anellovirus k-mer hits (119,599/142,612; Figure 1B). Strikingly, 76 of the 94 anellovirus k-mer-rich HCL runs were derived from the SKNO-1 cell line, an FAB-M2 t(8;21) AML cell line established in 1990 from the bone marrow of a 22-year-old man (25), accounting for 86.9% (103,988/119,599) of all anellovirus k-mer hits in these samples. The remaining anellovirus-rich HCL runs came from other human cell lines, including human pancreatic cancer PANC-1/HPAC (5 runs, 4,780 hits), primary neutrophils (4 runs, 3,854 hits), HepG2 (2 runs, 3,350 hits), BT549 (2 runs, 2,333 hits), K562 (1 run, 394 hits), HeLa (1 run, 390 hit), unknown (1 run, 264 hits), HEK_293T (1 run, 141 hits), and RD/18 rhabdomyosarcoma (1 run, 105 hits) cell lines (Figure 1C). Of the 76 SKNO-1 sequencing runs with high anellovirus k-mer counts, 66 were derived from RNA-Seq libraries, representing 88.9% of the total SKNO-1 anellovirus k-mer hits (92,398/103,988), while the remining 10 were ChIP-Seq (Chromatin Immunoprecipitation followed by sequencing) libraries, contributing 11.1% of SKNO-1 anellovirus k-mer hits (11,590/103,988; Figure 1D).

**Figure 1.**
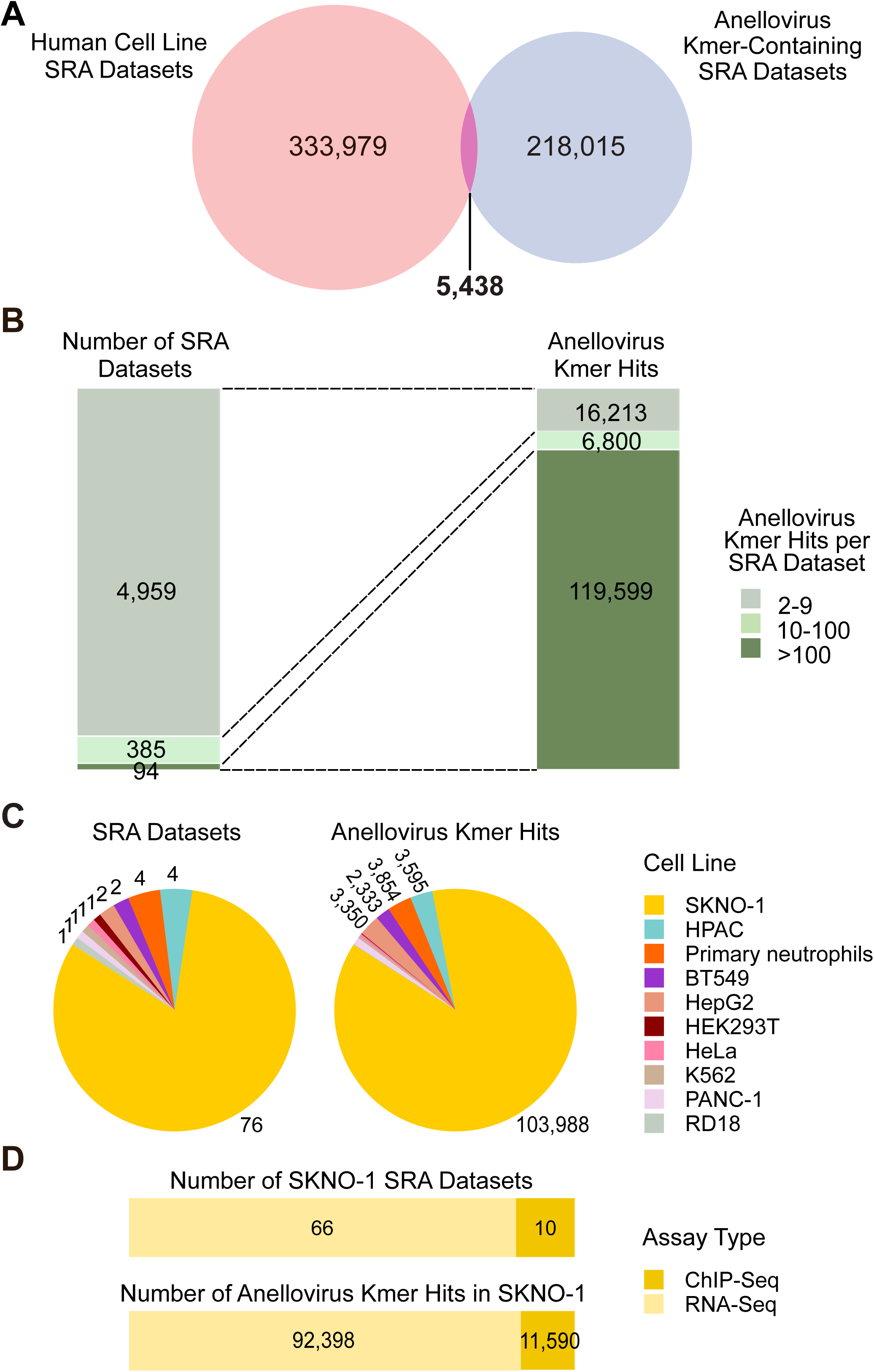
Data mining of NCBI SRA datasets for human cell lines with anellovirus k-mer hits. A) Venn diagram illustrating the overlap between human cell line SRA datasets and those containing anellovirus k-mers. B) Correlation analysis of SRA datasets with anellovirus k-mers hits, categorized by hit frequency: low (2-9 k-mers), medium (10-100 k-mers) and high (>100 k-mers). The dotted line connects each dataset to its corresponding number of hits. C) Classification of anellovirus k-mer-enriched SRA datasets by human cell line type. D) Overview of sequencing methodology for anellovirus k-mer-rich SKNO-1 SRA datasets.

To assess the potential role of contamination in these findings, we examined the origins of these anellovirus-rich datasets. The majority of RNA-Seq runs (41 FASTQ files) were derived from the University of Miami, which studied the enzyme co-activator CARM1 in normal and malignant hematopoiesis (26) and the transcriptional modulator DPF2 in inflammation (27). Other sequencing data were generated by the University Medical Center Mainz (24 RNA-Seq, 4 ChIP-Seq runs), where SKNO-1 was used in studies on oncogene regulation during acute resistance to BET inhibitors (28). In addition, six ChIP-Seq runs originated from the Broad Institute, focusing on transcriptional regulation associated with KMT2A-rearranged acute myelogenous leukemia (29), and one RNA-Seq run from the University of Virginia comparing transcriptomic profiling of AML primary samples and cell lines (30).

Searching SRA Cloud for sequencing runs with more than 1 anellovirus k-mer that were also annotated with “SKNO” yielded 138 sequencing runs, including 93 RNA-Seq, 41 ChIP-Seq, 3 ATAC-Seq (Assay for Transposase-Accessible Chromatin using sequencing), and 1 NA-Seq run (nuclease accessibility sequencing) (Table S4). Our initial SRA screen suggested both anellovirus DNA and RNA were present in the SKNO-1 cell line across a variety of laboratories from around the world.

### Assembly of anellovirus sequence from SKNO-1 cell line short-read sequencing

To create initial assemblies of anellovirus sequences from the SKNO-1 cell line given there were no classical DNA-Seq runs available for SKNO-1, we used four different RNA-seq runs of SKNO-1 cells with the highest number of anellovirus k-mers (Table 1). Specifically, we used a strand-specific RNA-Seq run of SKNO-1 cells transduced with a shRNA-Scramble from the University of Miami in 2018 (5,310 k-mers, SRA accession SRR6008465); an RNA-Seq run of BETiIC90-resistant SKNO-1 cells after 24h DMSO treatment from University Medical Center Mainz in 2022 (1,961 k-mers, SRR22306486); and an RNA-Seq run of SKNO-1 cells from University of Virginia in 2024 (1,391 k-mers, SRR24085425) (26, 28, 30). Based on literature review, we also included an RNA-Seq run of SKNO-1 cells published by the German cell line repository at Leibniz-Institute DSMZ (ERR3003594) in 2019 that had 559 anellovirus k-mers but was not annotated as “cell line” and so failed to be included in our initial screen conducted above (31).

**Table 1.** Assembly of anellovirus genome from four SKNO-1 RNA-Seq SRA datasets.

|  |  |  |  |  |
| --- | --- | --- | --- | --- |
| SRA Accession Number | SRR6008465 | SRR22306486 | SRR24085425 | ERR3003594 |
| Instrument | NextSeq 500 | NovaSeq 6000 | NovaSeq 6000 | HiSeq 2500 |
| Submission Year | 2018 | 2022 | 2024 | 2019 |
| Submitter | University of Miami, USA | University Medical Center Mainz, Germany | University of Virginia, USA | Leibniz-Institute DSMZ, Germany |
| Reference | (26) | (28) | (30) | (31) |
| Sequenced fragments/Spots | 54,934,628 | 44,374,157 | 20,228,673 | 34,619,305 |
| Read format | 2 x 75 bp | 2 x 150 bp | 2 x 151bp | 2 x 151bp |
| STAT anellovirus k-mers | 5,310 | 1,961 | 1,392 | 559 |
| CZID anellovirus reads (rpm) filecount | 14,295 (130.1) | 3,448 (38.8) | 4,095 (101.2) | 1,490 (21.5) |
| Longest de novo contig length | 3,322 bp | 2,976 bp | 3,375 bp | 3,331 bp |

Abbreviation: rpm, reads per million.

To ensure specificity, we used the CZID metagenomic pipeline to identify anellovirus-aligning reads (32). Across the four samples, 93.5 - 97.3% of reads were classified as human, with the only non-human viral genus being Anellovirus, except for Lentivirus in SRR6008465 which was used as vector for shRNA-Scramble transduction. We performed de novo assembly of the anellovirus-specific reads, yielding one to three contigs per dataset. The longest contigs ranged in length from 2,976 to 3,375 bp, aligning with 100% nucleotide identity over a 2,897 bp core region. Upon inspecting the alignment of the longest contigs, we noted that the RNA-Seq assembly from DSMZ dataset yielded a perfect 77 bp overlap between the ends of the longest contig, suggesting a circular viral genome. We trimmed the 5′-end 77 nt overlap and used the resulting 3,254 bp assembly as a reference genome to map reads from the other RNA-Seq datasets analyzed. Mapping the reads from four separate RNA-Seq runs showed that 99.7-100% of the reads classified as anellovirus perfectly aligned to the reference genome, with no significant nucleotide variability. This finding strongly suggests that there was no viral diversity in the analyzed samples, and a single anellovirus strain was present.

Interestingly, at the 5′ end of the alignment, we observed two distinct populations of reads: one aligned fully to the anellovirus genome, while the other was chimeric, showing a mix of human 28S rDNA gene (human chromosome 21, CT476837.18) at the 5′ end, and anellovirus sequence at the 3′ end (Supplementary Figure 1). At the 3′ end of the genome alignment, all anellovirus reads showed identity with the reference sequence but included soft-clipped sequences ranging from 30 to 92 nt that resulted to be identical to the 5′ end of the reference genome, consistent with a circular genome. In addition, we identified two single nucleotide variants (SNVs) in the assembly at positions 150 (c150a) and 162 (g162a), which were associated with reads that also displayed a 3′ end soft-clipped sequence extending up to 90 nt beginning at nucleotide 172 of the anellovirus genome (Supplementary Figure 1). These soft-clipped sequences showed 100% identity with an LTR-retrotransposon sequence in human chromosome 4 (AC107058.4), suggesting the presence of more chimeric sequences with human genomic material.

These findings were consistent across all four RNA-Seq runs, supporting the hypothesis that an integration event could be responsible for the observed chimerism. We also noted a coverage drop beginning at nucleotide 172 of the viral genome, which was associated with the chimeric reads aligning to human genomic sequences in this region. Finally, we examined RNA-Seq data from other non-SKNO-1 cell lines with anellovirus reads (Figure 1C), aligning them against both the human genome as well as their longest anellovirus de novo contig and searching for virus-human chimeric reads. While we confirmed the presence of different anelloviruses in these datasets, we did not observe the same pattern of virus-human chimeric reads, suggesting that this feature is specific to the anellovirus in the SKNO-1 cell line.

### Long-read sequencing of the SKNO-1 anellovirus integrated into human chromosome 21

To confirm the potential integration of an anellovirus in the SKNO-1 cell line, we obtained SKNO-1 cells from the DSMZ repository in Germany (G subline) and the JCRB repository in Japan (J subline) and performed 40x PacBio sequencing of high-molecular weight dsDNA isolated from these cell lines. We obtained 8,683,919 reads with an N50 length of 16,563 bp from the SKNO-1 G cell subline, and 8,366,596 reads with an N50 length of 18,656 bp from the SKNO-1 J cell subline. Mapping these PacBio reads to the draft assembled anellovirus genome from DSMZ RNA-Seq revealed five reads (median length 17,024 bp) for the G subline and 12 reads (median length 18,957 bp) for the J subline. Aligning the anellovirus-mapping PacBio reads to each other recovered a 3,425 bp anellovirus sequence that consisted of the 3,254 bp draft DSMZ anellovirus genome assembled from RNA-seq reads followed by a repeat of the 5′ end 171 bp of the contig at the 3′ end (Figure 2A-B). These repeats contained the two nucleotide variations – c150a and g162a – which were previously identified in our RNA-Seq assembly.

**Figure 2.**
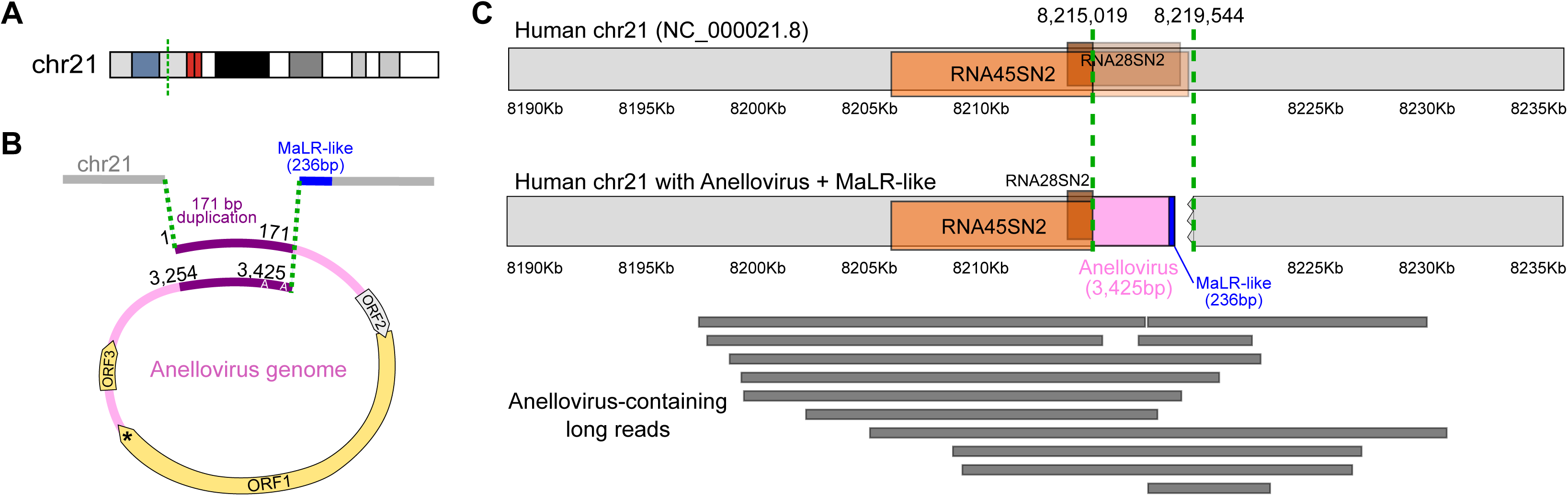
Integration of the anellovirus genome within chromosome 21 of the SKNO-1 cell line. A) Schematic representation of chromosome 21 in the SKNO-1 cell line, showing the integration site of the anellovirus genome (dotted green line). Red represents the centromere, blue indicates the repetitive region, and the grayscale sharing indicates Giemsa bands (darker shades correspond to higher heterochromatin and greater AT-rich regions). B) Detailed schematic of anellovirus integration within chromosome 21. The pink region denotes the anellovirus genome, with arrows indicating open reading frames (ORF). A grey arrow highlights ORF2 containing an early stop codon at position codon position 2, while an asterisk (*) marks the truncation of ORF1, preventing overlap with ORF3. The violet line highlights the duplicated sequences located at both termini of the viral genome, which differ by two-point mutations (C→A and G→A), while the blue region indicates an additional nucleotide segment corresponding to a MaLR-like element. C) Comparison between wild-type chromosome 12 and the chromosome 21 with the anellovirus genome integration. The integrant interrupts the RNA45SN2/RNA28SN2 rRNA gene. Dotted green lines indicate the region that has been altered within chromosome 21 including specific coordinates for the integration site. Below the schematic of chromosome 21 with the integration, PacBio long reads are shown (represented by grey rectangles), confirming the presence of the integrant in the SKNO-1 J subline cell.

Further analysis revealed that the anellovirus genome was flanked by human sequences. NCBI BLAST searches identified these sequences as being part of human chromosome 21, specifically within the rDNA genes *RNA45SN2* and *RNA28SN2* (Figure 2C). The full integration region spanned from 8,215,019 to 8,219,545 on human chromosome 21 (NC_000021.9). The integrated sequence consisted of the 3,425 bp anellovirus genome, followed by 10 bp at the 3′ anellovirus-human junction that could not be further characterized, and 146 bp that perfectly matched the negative strand of human chromosome 4 (coordinates 65,017,569 - 65,017,714, NC_000004.12), with 82 bp that could not be aligned to anything in the nt/nr database by BLASTn. Overall, the integration replaced 4,522 nt of human sequence with a 3,661 bp integrant.

The structure of the integrated viral genome was supported by contiguous read alignments extending up to 17,719 bases at the 5′ end and 11,263 bases at the 3′ end of the viral sequence (Figure 2C). Notably, the integration junctions were identical across both SKNO-1 G and J cell sublines. The 5′ junction occurred at a homopolymeric stretch of 8 guanosines within chromosome 21, resulting in the loss of one human G-quadruplex motif. This event left a single G-quadruplex element at the beginning of the integrated genome, followed by the 171 bp terminal repeat at the 3′ end. In addition, two of the 12 anellovirus-containing reads from the SKNO-1 J subline extended over 12 kb upstream of the integration site on human chromosome 21 and captured only the first 308 nucleotides of the anellovirus integrant and then continued for 2,592 nt on chromosome 21, thus omitting the rest of the anellovirus genome and the chromosome 4-like sequence.

To further explore the integration, we analyzed the depth of sequencing coverage for the anellovirus and several housekeeping genes (Supplementary Figure 2). In both G and J sublines, we observed overall consistent chromosomes coverage with slight variations that reflect the underlying karyotypic heterogeneity of SKNO-1 cell line (Supplementary Table 5). Specifically, *SDHA* (chr5) showed elevated depth (∼68X), consistent with the known gain of chromosome 5 in the SKNO-1 karyotype. *HPRT1* (chrX) showed depth values (23.2x in G, 15.7x in J) aligned with expectations for a single-copy X chromosome in a male cell line. *SOD1* (chr21), located near the anellovirus integration site, showed slightly reduced depth in the J line (31.6x) relative to G (40.0x). Intriguingly, the depth of anellovirus coverage (8.3x in J vs. 4.7x in G) was less than expected single-copy coverage in this clonally heterogeneous cell line (25).

### Quantification of anellovirus in SKNO-1 cell line with ddPCR

To quantify Alphatorquevirus sp. isolate within SKNO-1 cell line, we performed ddPCR with two primer/probe sets. We used primer/probe set 1 (PP1), designed to target the negative strand within the ORF1 coding sequence, and primer/probe set 2 (PP2), targeting the positive strand in the 171-bp duplicated ends corresponding to non-coding sequence. The virus copy number was quantified in SKNO-1 J and G sublines, with THP-1 monocytes and 293A kidney cells serving as negative controls. Using *RPP30* as the reference gene (assuming 2 copies/cell based on PacBio data, Figure S2), the anellovirus copy number was 0.54-0.56 copies/cell in J subline and 0.42-0.53 copies/cell in the G subline. No anellovirus was detected in THP-1 or 293A cells.

Next, we verified RNA expression of Alphatorquevirus sp. in SKNO-1 J and G sublines using RT-ddPCR with the same primer/probe sets and negative controls as ddPCR. Transcripts were detected in both J and G sublines. Relative RNA expression, measured as PP1(ORF1):RPP30, was 7.5 in J subline and 9.0 in G subline, while PP2(non-coding):RPP30 expression was 3.4 in J subline and 3.8 in G subline. These results suggest that viral transcripts are generated from ORF1, and that a lower level of transcription also occurs from the non-coding viral sequence within the rDNA locus. No significant DNA contamination was observed in the RNA samples. No anellovirus transcripts were detected in THP-1 or 293A cells.

### ChIP-Seq and RNA-Seq Data Analysis of Integrated Anellovirus

Next, we used the assembly of viral-host integrant sequence from PacBio long read sequencing as a reference for analyzing a variety of SKNO-1 RNA-Seq (Supplementary Figures 3 and 4) and ChIP-Seq datasets (Figure 3 and Supplementary Figure 5). RNA-Seq data showed sequencing coverage across the integrant sequence including the viral ORFs and intergenic regions. Assemblies also showed reads supporting canonical splicing of SKNO-1 anellovirus mRNAs in ORF2 to create ORF2/2 and ORF2/3 (Supplementary Figure 3).

**Figure 3.**
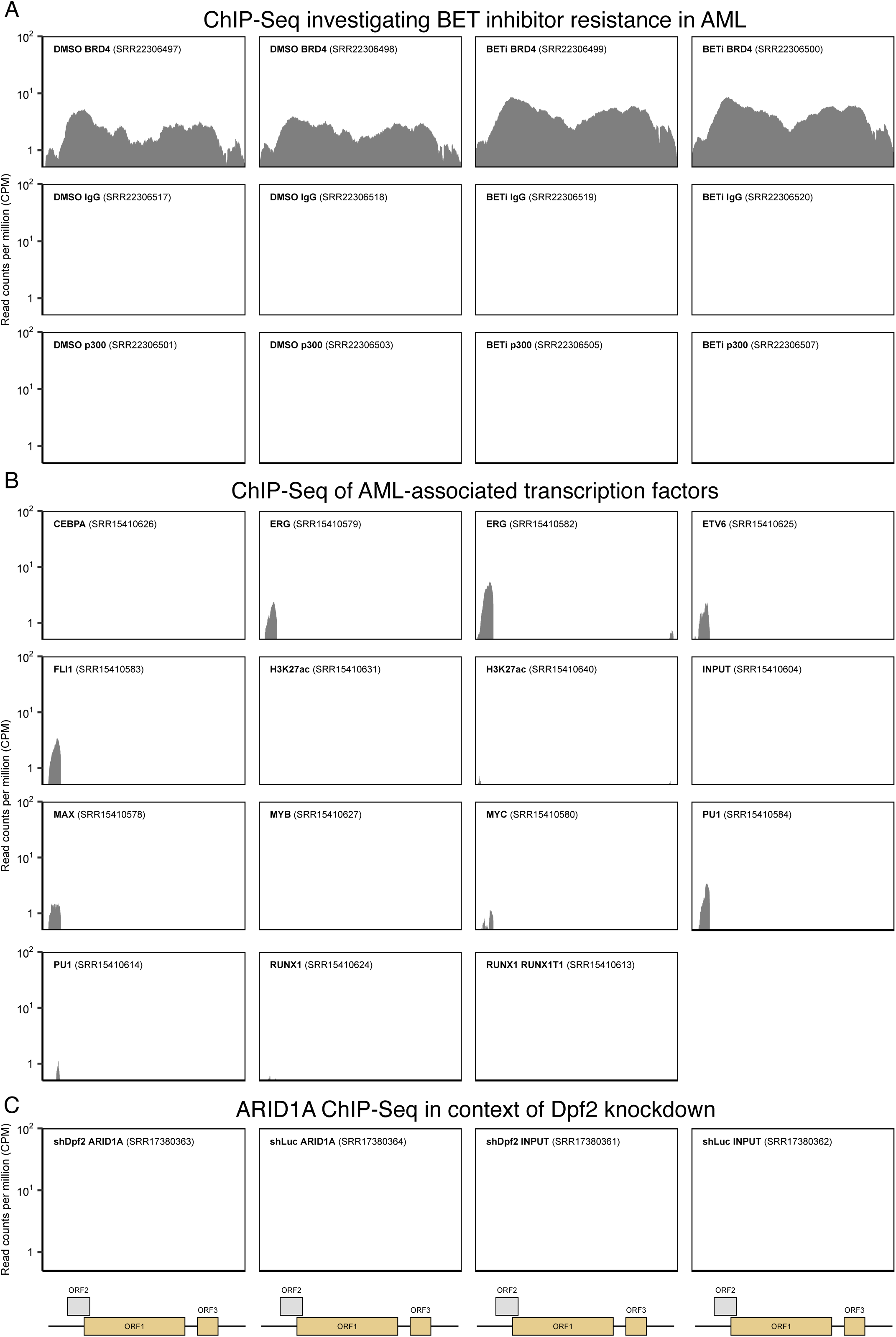
ChIP-Seq SRA data mapping to anellovirus SKNO-1 assembly. Reads from ChIP-Seq SRA datasets were trimmed with fastp and aligned to a de-novo assembly of the anellovirus genome constructed from RNA-Seq data of the SKNO-1 cell line (see Methods). Alignment depth is plotted across the 3,425 bp reference genome. Plots are labeled with the associated SRA and ChIP-Seq experimental conditions (e.g. protein targeted, buffer). Predicted open reading frame coordinate intervals of the anellovirus assembly are displayed at the bottom. The span of the entire assembly is depicted by the black bar, and ORFs are shown as yellow boxes.

ChIP-seq data from studies using SKNO-1 cells, which examined the response to BET protein inhibition, showed that BRD4, a bromodomain-containing protein that canonically facilitates transcription by recruiting transcriptional machinery to acetylated chromatin, binds broadly across nearly the entire Alphatorquevirus genome (Figure 3) (28). Notably, BRD4 occupancy at the viral integration site persisted and even slightly increased following BET inhibition, indicating partial or full resistance to bromodomain inhibition, while p300 chromatin occupancy was minimal. In addition, treatment with BET and p300 inhibitors and selection of drug-resistant SKNO-1 cells increased overall anellovirus transcription by between 2-4-fold (Supplementary Figure 4). This suggests either noncanonical recruitment of BRD4 to the anellovirus locus or maintenance within a stable, enhancer-like chromatin domain, consistent with an active and accessible regulatory region.

Focal enrichment was observed for hematopoietic transcription factors MAX and the ETS family (ERG, FLI1, SPI-1/PU.1, and ETV6) within the ∼300 bp upstream of the viral open reading frames (Figure 3) (29). The two tagging SNPs in the 171 bp repeat region (c150 and g162) demonstrated only the 5′ repeat was recovered in the transcription factor ChIP-seq reads. High-scoring motifs were found in the first 312 bp of the viral integrant for human SPI-1/PU.1, MAX, ETV6, and FLI-1 transcription factors using FIMO v5.5.9 analysis from the MEME suite (p<0.0001, Supplementary Table 6) (33), while the ERG motif had >0.88 relative profile score using JASPAR2026 (Supplementary Table 7) (34). Interestingly, no H3K27ac enrichment was observed, supporting the noncanonical BRD4 recruitment described above. These data indicate that the integrated viral DNA contains discrete regulatory elements that result in transcription factor binding within a broadly accessible BRD4-enriched chromatin domain.

### Phylogenetic analysis and characterization of integrated SKNO-1 Alphatorquevirus

The SKNO-1 anellovirus genome, integrated into chromosome 21, is 3,425 bp in length with a GC content of 50.6%. BLAST analysis identified *Alphatorquevirus homin 29* as the closest known viral species, sharing 73.6% nucleotide identity with the ICTV reference genome Torque teno virus 29 (AB038621.1, length 3,676 bp). Notably, the first 422 nucleotides of the Torque teno virus 29 genome were absent in the SKNO-1 anellovirus, and no homologous sequence were found in the SKNO-1 anellovirus genome by dot plot analysis. Interestingly, the last 316 nt of the Torque teno virus 29 genome are homologous to the first 316 nucleotides of the SKNO-1 anellovirus integrant.

We predicted the ORF1 of the SKNO-1 anellovirus using an alternative ACG start codon, based on homology with Torque teno virus 29, yielding 80.8% nucleotide identity and 88.9% amino acid similarity (BLOSUM62). SKNO-1 ORF1 contains an early stop codon at Trp565*, producing a protein truncated by 171 amino acids relative to the homolog. The predicted ORF3 shows 87.5% nucleotide identity and 82.6% amino acid similarity. In contrast, the SKNO-1 anellovirus ORF2 contained a mutated start codon (atg > act), a stop codon at amino acid position 2, and had limited homology in the first 18 nt with the Torque teno virus 29 ORF2, suggesting that SKNO-1 anellovirus ORF2 is either not expressed or may initiate translation from an in-frame alternative CTG start at codon 7 of the Torque teno virus 29 ORF2.

To confirm the genus and species classification of the SKNO-1 anellovirus, we performed pairwise comparison and phylogenetic analysis of ORF1 sequences from 92 reference genomes across the Alpha-, Beta-, and Gammatorquevirus genera (Figure 4). The phylogenetic tree confirmed that the SKNO-1 anellovirus clusters within the *Alphatorquevirus* genus. Pairwise nucleotide comparison showed that SKNO-1 ORF1 exceeds the 69% nucleotide identity threshold relative to *Alphatorquevirus homin 29* (AB038621.1), either individually or within its closest evolutionary clade, supporting its classification as a species of *Alphatorquevirus homin 29* (28).

**Figure 4.**
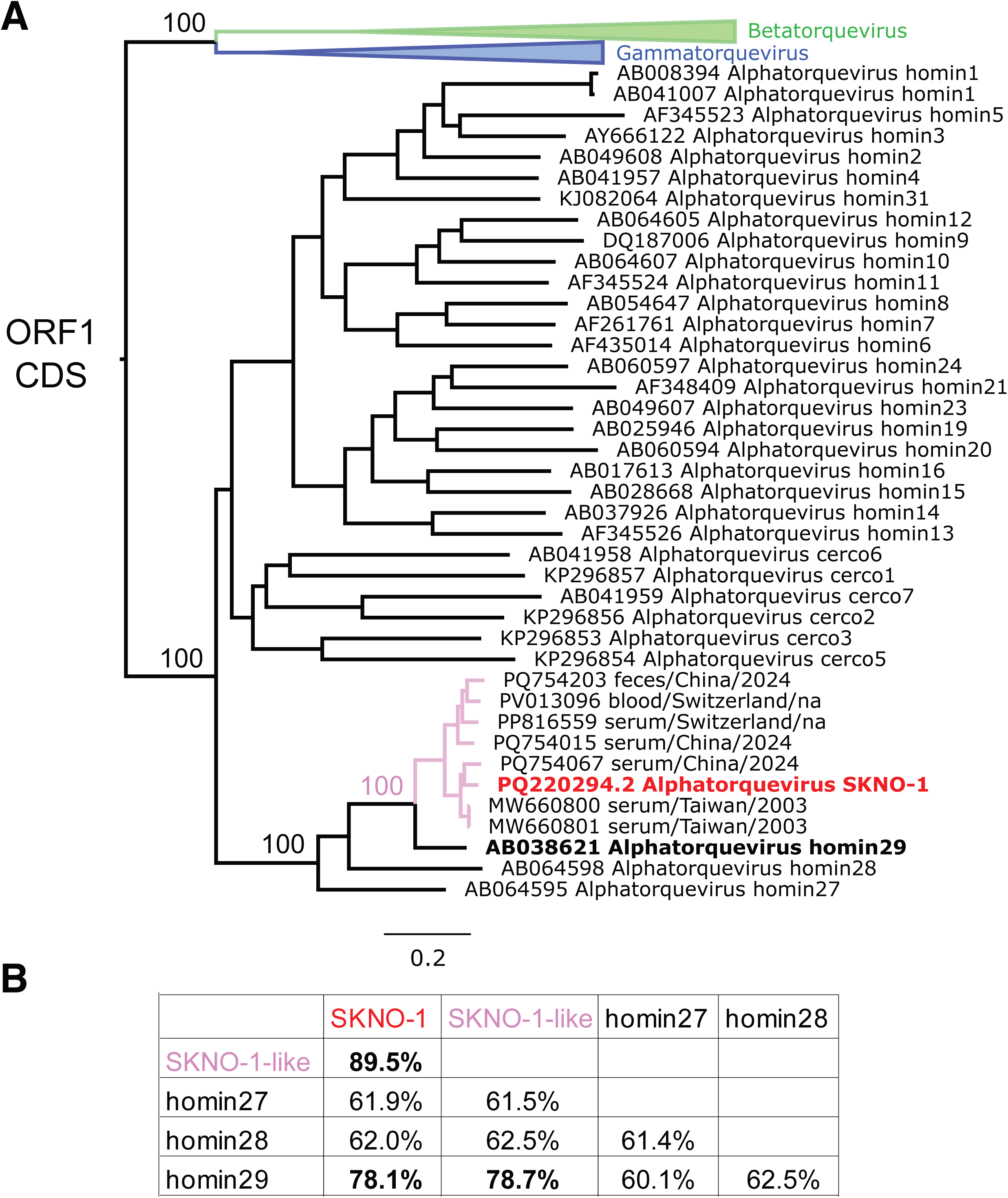
Phylogenetic analysis of anellovirus. A) Maximum-likelihood phylogenetic tree of ORF1 nucleotide sequences from Alphatorquevirus, Betatorquevirus, and Gammatorquevirus reference genomes. Ultrafast bootstrap support values (1,000 replicates) are shown for key nodes. Anellovirus assembled from the SKNO-1 cell line is highlighted in red, and branches corresponding to the clade containing this virus together with sequences retrieved from NCBI BLAST showing >90% genomic identity are shown in pink. The Betatorquevirus clade is collapsed and represented as a green triangle, while the Gammatorquevirus clade is collapsed in blue. Branch lengths correspond to the number of substitutions per site. B) P-distance matrix comparing the SKNO-1 anellovirus, the SKNO-1–like anellovirus clade, and representative sequences of Alphatorquevirus homin27, homin28, and homin29. Bootstrap standard errors (100 replicates) are shown in italics.

ORF1 encodes the longest viral protein and functions as the capsid protein (36). In SKNO-1, ORF1 is predicted to procude a 564-amino acid (aa) protein initiating from an ACG start codon (37), considerably shorter than the average Alphatorquevirus ORF1 (749 aa), and even shorter than the Betatorquevirus ORF1 (664 aa). Due to lack of experimental validation for the homology-predicted ACG start codon in this Alphatorquevirus clade, we also considered an alternative ORF1 prediction of 476 aa initiating at an in-frame ATG codon.

To investigate the structural features of SKNO-1 ORF1, we performed homology modeling using the only published anellovirus ORF1 structure (Betatorquevirus LY1 ORF1, PDB: 8V7X) (36). Alignment indicated that SKNO-1 ORF1 retains the jelly roll (JR), P1, and P2 domains, but lacks the C-terminal region (Figure 5A). Since the LY1 ORF1 structure was determined without the ARM or C-terminus, the truncated C-terminus in SKNO-1 ORF1 is not expected to substantially alter the predicted protomeric structure or a putative pentamer, as confirmed by AlphaFold3 modeling (Figure 5B). Electrostatic surface analysis further indicates a similar charge distribution between LY1 ORF1 and SKNO-1 ORF1 (Supplementary Figure 6), suggesting that biophysical properties are largely conserved despite truncation.

**Figure 5.**
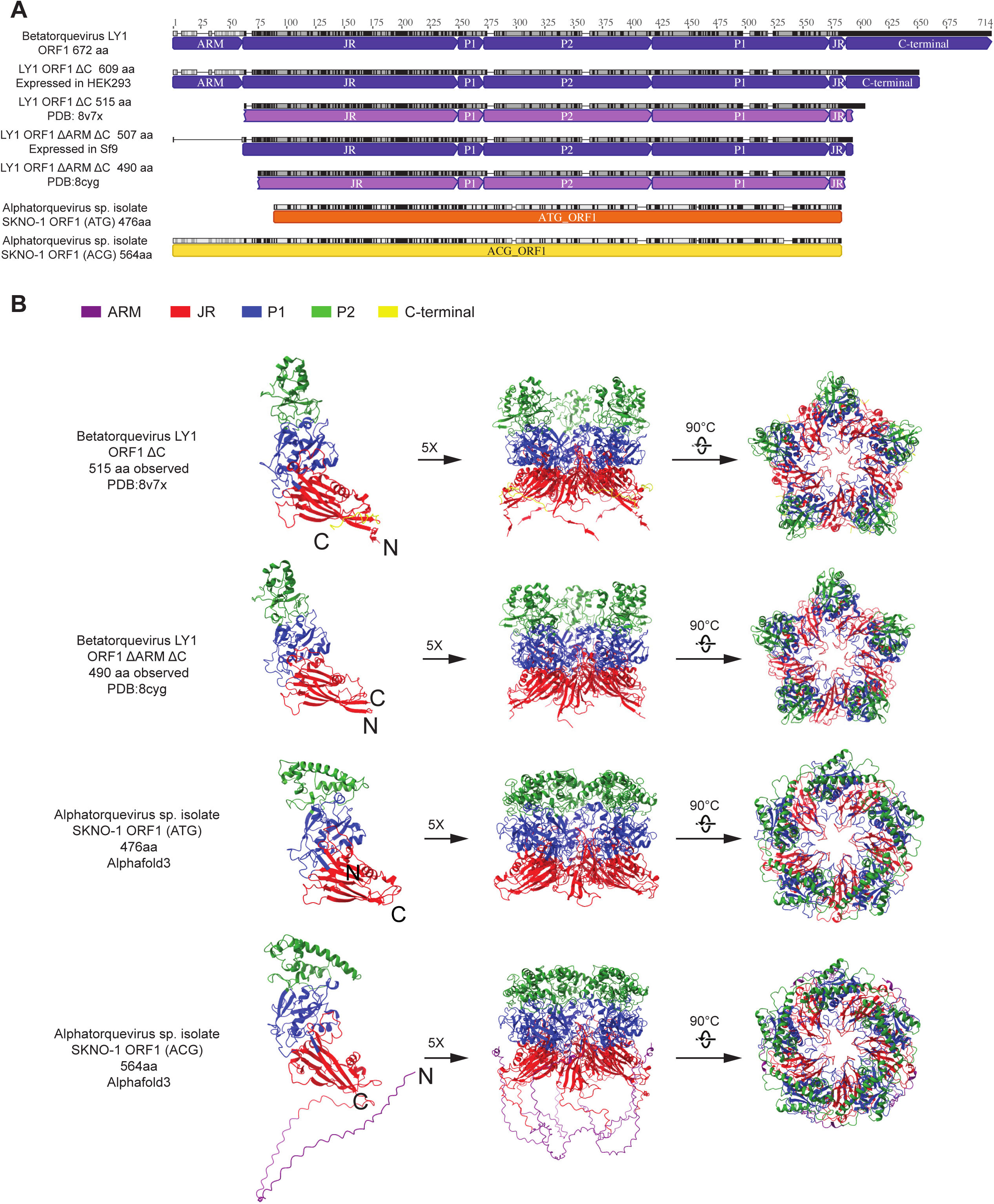
Alphatorquevirus sp. isolate SKNO-1 ORF1 (capsid protein) analysis. A) Multiple alignment of amino acid sequence between Betatorquevirus LY1 ORF1 and Alphatorquevirus sp. isolate SKNO-1 ORF1 starting with ATG or ACG. The identical sites are marked as black. Betatorquevirus LY1 ORF1 is annotated with purple bar, and Alphatorquevirus sp. isolate SKNO-1 ORF1 is annotated with orange and yellow bar. B) The PDB structures of Betatorquevirus LY1 ORF1(33) and AlphaFold predicted structure Alphatorquevirus sp. isolate SKNO-1 ORF1 starting with ATG or ACG. The motif ARM, P2, P1, JR, and C-terminal are marked on the structure. Pentamers structures are shown on the right.

## DISCUSSION

Here, using a combination of large-scale in silico screening, long-read sequencing, and molecular validation, we identified a stable integration of an Alphatorquevirus homin29 genome within the rDNA locus of the AML cell line SKNO-1 (25). The integration replaces a 4.5 kb segment of RNA45SN2/RNA28SN2 on chromosome 21 with a 3.6 kb insert containing a complete viral genome, a short unassigned junction, a MaLR-like retrotransposon fragment, and an unclassified sequence. The recurrence of this integration in independently sourced SKNO-1 lines supports that this is a genuine event rather than contamination. Quantification by ddPCR indicates ∼0.5 viral genomes per cell, consistent with a subclonal representation, while RT-ddPCR confirms transcription of ORF1 as well as the non-coding viral terminal repeats, revealing that the complete integrated element as well as the viral ORF1 are transcriptionally competent. We detected spliced ORF2/2 and ORF2/3 transcripts despite an early stop codon observed in the predicted ORF2 sequence (10). Structural modeling suggests that truncated ORF1 retains key capsid domains and physicochemical properties, hinting at preserved structural potential despite probable loss of full virion production (36).

Our data raise a number of intriguing hypotheses around anellovirus biology in humans. The existence of a fossilized anellovirus integrant in a t(8;21) AML cell line – thought to be derived from an early myeloid progenitor cell (38, 39) – suggests that the anellovirus was present as an episomal genome in the same early myeloid progenitor at the time of leukemia initiation. Integrated anellovirus DNA have recently been found to be a driver of acute promyelocytic leukemia (40–42), whose leukemia-initiating promyelocyte is one step developmentally downstream of the early myeloid progenitor. It has long been recognized that anellovirus levels are highest in peripheral blood and bone marrow and can be detected in both mature lymphoid and myeloid cells (3, 43–46). A simple model for the high prevalence of anelloviruses in blood and blood cells would be residency within hematopoietic stem cells, and our data suggest a cell-of-origin that is developmentally one step closer to the long-term hematopoietic stem cell progenitor. Alternatively, anelloviruses may simply have the ability to infect any cell due to lack of a specific receptor. Certainly, more work is needed to understand mechanisms of anellovirus entry.

To date, relatively little is known about the mechanisms that govern anellovirus transcription. Suzuki et al. (2004) and Kamada et al. (2004) both mapped promoter and enhancer regions within the upstream untranslated region of Alphatorqueviruses, and Suzuki et al. (2004) showed upstream stimulating factor (USF) family proteins bound TTV promoter and removal of the USF binding site was associated with reduced promoter activity (47, 48). Since 2004, thousands of new human anellovirus sequences have been recovered and it is unclear whether these mechanisms are conserved among all anelloviruses. For instance, while the TATA box sequence (TATAA) is present in the same location in Alphatorquevirus homin 27, 28, and 29 type species (and mutated to TATTA in SKNO-1 anellovirus), the USF binding site is wholly absent in Alphatorquevirus homin 27, 28, and 29, nor is it present in the SKNO-1 anellovirus. Through data mining of cancer biology-funded systems biology data of AML cell lines, we offer new hypotheses around hematopoietic cell transcription factors such as ETS family members that can specifically bind anellovirus untranslated region sequence and are associated with viral gene transcription. Future work can experimentally deplete these transcription factors to examine their requirement for anellovirus transcription.

The mechanism by which the SKNO-1 anellovirus integrated into the human genome is a matter of speculation. The integration may have arisen via GGGG microhomology-mediated end joining with template switching, a mechanism analogous to those described for hepatitis B virus, human herpesvirus 6A, and human papillomaviruses (49–52). While alternative scenarios – such as prior MaLR insertion or non-homologous end joining followed by recombination – cannot be excluded, they are less parsimonious. It is also unclear which RNA polymerase is responsible for the anelloviruses transcription. The viral sequence integrated in a transcriptionally permissive locus within an RNA Pol I-transcribed rDNA unit. This region exhibits broad BRD4 occupancy and focal binding by hematopoietic transcription factors MAX and ETS (ERG, FLI1, PU.1, ETV6), which are typically associated with RNA Pol II transcription. Persistent BRD4 engagement despite BET inhibition suggests bromodomain-independent tethering, potentially with coactivators retained likely through protein–protein interactions rather than high histone acetylation density, consistent with absent H3K27ac ChIP-Seq enrichment (53). Future work involving ChIP-Seq of RNA Pol occupancy or nascent RNA labeling is required to understand whether these features create an “RNA Pol II island” within the nucleolar organizer region, where local chromatin remodeling and sequence-specific elements facilitate transcription of the integrated viral genome, in line with RT-ddPCR detection of both ORF1 and the non-coding viral region.

The presence of the anellovirus integrant in different sourced SKNO-1 and high similarity (>90%) to other Alphatorquevirus genomes detected in human clinical samples (collected in 2003 and 2024) suggests it occurred in vivo during leukemogenesis and represents a contemporary viral variant. The subclonal nature of the integrant and its long-term persistence in culture suggest cellular tolerance rather than a selective advantage. Although anelloviruses are typically non-integrative, sporadic integration events have been described, and recent reports suggest that host DNA repair pathways can occasionally capture episomal anellovirus DNA (42). These observations support a model in which the SKNO-1 integrant represents a stochastic, repair-mediated event rather than a biologically selected insertion. Thus, this case likely represents a rare example of somatic viral capture with minimal phenotypic impact, mechanistically analogous, but not functionally comparable, to the repair-driven integrations seen with other non-integrative DNA viruses (49, 51).

This study has a number of limitations. While SKNO-1 provides a striking experimental case of viral integration into human DNA, it does not recapitulate natural anellovirus infection biology in humans, where the virus exists as an episome and persistence would not require integration. While transcriptional activity of the anellovirus locus is evident, production of protein or infectious virions remains unconfirmed. Of note, we attempted to assay for protein production, raising rabbit polyclonal antibodies to 18 and 20 aa peptides in ORF1 P1 and P2 regions, respectively. The anti-P1 antibody showed no signal in Western Blot at the expected ORF1 size, while the anti-P2 antibody gave a band of expected size in SKNO-1 lysate but this was band was also non-specifically present in HEK293 cells and we could not recover anellovirus peptides by immunoprecipitation mass spectrometry with the anti-P2 antibody. Our inability to recover protein could be due to the truncated nature of ORF1, although removing the C-terminal region was helpful for stable ORF1 production in insect cells and to prevent proteolysis (36). To date, no one has seen bona fide ORF1 protein from Alphatorquevirus species annotated with an alternative ACG start codon that includes the conserved N-terminal arginine rich motif. Future work could use ribosome profiling of SKNO-1 cells and/or these Alphatorquevirus species to determine ribosomal occupancy of anellovirus transcripts and whether ORF1 protein is produced. As stated above, we do not know if the ETS family transcription factors are required for SKNO-1 anellovirus transcription, which would require specific depletion and examination of SKNO-1 anellovirus expression. Finally, we also do not definitively know if the anellovirus was present in the leukemia-initiating cell for this t(8;21) AML, generally thought to be an early myeloid progenitor. Though less likely in our estimation, alternative models could be the anellovirus infected the cells that would become the SKNO-1 line after leukemogenesis or early during cell line establishment. Future work should examine single-cell RNA-sequencing data to examine the tropism of anelloviruses in bone marrow cells and throughout the human body.

In summary, this study highlights the power of public sequencing data as well as the plasticity of anellovirus genomes and their capacity to integrate in transcriptionally active host loci. The SKNO-1 integration demonstrates how anelloviruses can be tolerated by host cells over long-term culture, offering a unique model to explore the interplay between viral transcription, chromatin environment, and host regulatory factors of an understudied virus. This system provides a foundation for future studies on anellovirus transcriptional regulation, epigenetic interactions, and potential impacts on hematopoietic biology.

## MATERIALS AND METHODS

### Data mining in NCBI SRA cloud by BigQuery

NCBI SRA accession numbers containing reads taxonomically classified as *Anellovirus* (taxid: 687329) were identified using the NCBI SRA Taxonomic Analysis Tool on BigQuery (https://www.ncbi.nlm.nih.gov/sra/docs/sra-cloud/). This dataset is based on k-mer-based classification, typically using 32-nt k-mers, performed by NCBI. The list of accessions was obtained on July 4, 2025, using the following command: bq --format=csv query --nouse_legacy_sql --max_rows=10000000 ‘SELECT acc, total_count FROM ‘nih-sra-datastore.sra_tax_analysis_tool.tax_analysis’ WHERE tax_id = 687329 and total_count > 1’.

Separately, NCBI SRA runs annotated as derived from human cell lines were retrieved on July 4, 2025 using the following query: bq --format=csv query --nouse_legacy_sql --max_rows=1000000 ‘SELECT acc, assay_type, jattr, releasedate FROM ‘nih-sra-datastore.sra.metadatà WHERE organism = “Homo sapiens” and contains_substr(attributes, “cell line”)’.

We then identified the subset of human cell line runs that also contained Anellovirus k-mer evidence, by matching accession numbers across both datasets. These overlapping runs represent cases where anellovirus was detected in human cell line sequencing data, despite not being explicitly reported in the sample metadata. The matching accessions were retrieved with the following command: bq --format csv query --nouse_legacy_sql --max_rows=10000000 ‘SELECT t1.acc as ‘Accession Number SRÀ, total_count as ‘Anellovirus K-mer Count’, assay_type as ‘Assay Typè, TO_JSON_STRING(jattr) as attributes FROM (SELECT * FROM ‘nih-sra-datastore.sra_tax_analysis_tool.tax_analysis’ WHERE tax_id = 687329 and total_count > 1) AS t1 INNER JOIN (SELECT * FROM ‘nih-sra-datastore.sra.metadatà WHERE organism = “Homo sapiens” AND contains_substr(attributes, “cell line”)) as t2 ON t1.acc = t2.acc. Accession lists for each dataset are provided in Tables S1–S3.

### SKNO-1 RNA-Seq data analysis

From the list of SKNO-1 cell line sequencing containing anellovirus reads, we downloaded three RNA-seq FASTQs with the highest anellovirus-kmer counts, originating from different laboratories (NCBI SRA accession numbers SRR22306486, SRR24085425, and SRR6008465 (Table 1). In addition, RNA-seq dataset ERR3003594 was analyzed, as it corresponds to the dataset characterizing the SKNO-1 cell line strain obtained from DSMZ repository that was further utilized in this study. Metagenomic analysis was performed by CZID cloud-based metagenomic platform accessed on July 05, 2025 (32). From each FASTQ file, reads classified as family *Anelloviridae* in CZID were downloaded and de novo assembled using SPAdes v4.2.0 with the “isolate” mode and PHRED score offset 33. The longest contig of each de novo assembly was chosen for NCBI BLASTn using the non-redundant nucleotide (nt/nr) database. We further evaluated the de novo assemblies by mapping the FASTQ reads against the longest obtained contig using minimap2 (version 2.29-r1283) with parameters to adjust for short read RNA-seq, considering splicing and discarding secondary alignments.

To search for human-anellovirus chimeric reads in SRAs obtained from other cell lines, we first used Kraken2 (v2.17.1) to isolate anellovirus-classified reads (54). To construct the Kraken2 database used for classification, we downloaded all NCBI nucleotide entries under the *Anelloviridae* taxon (ID: 687329) with genome length between 2,000-8,000 bp (n=24,398 sequences) and filtered out entries with large homopolymer regions and/or interior sequence homologous to the human genome consistent with mis-assembly, resulting in 22,255 sequences. Filtering was performed with bbduk.sh from the BBTools suite (v39.26) with parameters maxns=0 entropy=0.955 entropyk=5 entropywindow=100 (55). Similarity to human sequence was manually assessed using BLASTN against *Homo sapiens.* Reads were then classified to a database containing these sequences via Kraken2 with parameters --minimum-hit-groups 2. Unclassified reads were discarded and classified reads were aligned to human genome GCA_000001405.15_GRCh38 (comprised of chrs 1-23, X, and Y, sourced from the analysis set at https://hgdownload.gi.ucsc.edu/goldenPath/hg38/bigZips/analysisSet/) using minimap2 (v2.29-r1283) with parameters -L --secondary no -ax splice:sr and samtools view (v1.21) with parameters -F 2052 (56). As chimeric reads result in only partial alignments, mapped reads were filtered with a custom python script to have at least 20% of their read length comprised of soft-clipped, unaligned nucleotides. Reads surviving this filter were then realigned to the same *Anelloviridae* reference database used to construct the kraken2 database using minimap2 with the same parameters. To determine if lack of anellovirus alignment in the samples was due to a lack of coverage, we also de novo assembled anellovirus genomes from kraken2-classified reads using spades (v 4.2.0) with parameters --phred-offset 33. Code is available at https://github.com/greninger-lab/anellovirus_in_skno1_paper/.

### SKNO-1 ChIP-Seq data analysis

From the list of SKNO-1 sequencing containing anellovirus reads, we downloaded 10 ChIP-Seq FASTQ files from NCBI SRA. For control purposes, we also downloaded additional 29 ChIP-Seq SKNO-1 cell line FASTQ files from the same BioProjects but that did not contain anellovirus reads. As most of SKNO-1 ChIP-Seq runs are single-end (SE), we used only R1 reads of the pair-ended runs to keep consistency. Metagenomic analysis was performed as described in RNA-Seq analysis section. The reads coverage generated by minimap2 was then normalized by the total reads number to calculate reads per million (RPM). The sequencing depth profile was visualized in RStudio v2023.06.1 with ggplot2 and Adobe Illustrator 2022.

### Cell culture and collection

Two stocks of the acute myeloid leukemia SKNO-1 cell lines were procured from the DSMZ repository in Germany (G line) and the JCRB repository in Japan (J line). Cells were cultured in RPMI 1640 medium (Life Technologies Corporation, Carlsbad, USA) supplemented with 10% fetal calf serum (Life Technologies Corporation, Carlsbad, USA), and 10 ng/mL recombinant human GM-CSF (Life Technologies Corporation, Carlsbad, USA). THP-1 cells (ATCC, Manassas, USA) were cultured in RPMI 1640 medium supplemented with 10% fetal calf serum. 293A cells (ATCC, Manassas, USA) were grown in DMEM (Life Technologies Corporation, Carlsbad, USA) supplemented with 10% FBS. Cells were kept in a humidifier incubator at 37°C with 5% CO_2_.

### DNA and RNA extraction

Nucleic acid was extracted adding 0.1 mL chloroform to the cells in 0.5 mL TRI reagent. After vortexing and 5 min incubation at room temperature, samples were centrifuged at 12,000 x g at 4°C for 15 min. DNA was isolated from the interphase and the lower phenol-chloroform phase saved from RNA extraction. DNA was precipitated adding 0.15 mL of 100% ethanol. After washing twice with 0.1 M sodium citrate in 10% ethanol (pH 8.5) and once with 75% ethanol, DNA was resuspended in 8 mM NaOH. The insoluble materials were removed by centrifuging at 12,000 x g 4°C for 10 min. The pH of DNA solution was finally adjusted to 7.2 with 0.1M HEPES.

Carefully moving the aqueous phase into 0.25 mL isopropanol, RNA was precipitated at −80°C for an hour. The precipitated RNA was collected by centrifuging at 12,000 x g, 4°C for 10 min. After washing twice with 75% ethanol and air dry, 45 µL DNase/RNase-free water were added to resuspend the RNA. DNA was digested with 5 µL 10X Turbo DNase Buffer and 1 µl Turbo DNase from TURBO DNA-free Kit (Thermo Fisher Scientific, Vilnius, Lithuania). After incubating at 37°C for 30 min, the enzyme and divalent cations were removed by 5 µl DNase Inactivation Reagent to have DNA-free RNA.

### ddPCR and RT-ddPCR assay

Two primer/probe sets were designed using Primer3 based on the complete SKNO-1 anellovirus genome. The first set (PP1) targeted the ORF1 negative strand, with forward primer 5′-TGCCAATCAAGTGGTGGGTT-3′, reverse primer 5′-GAGGGTTTCGTCTGGTCTCG-3′, and probe 5′-/56-FAM/CAATAGTTA/ZEN/ACTGTGGACCGTTTGTCCCAC/3IABKFQ/ −3′. The second set (PP2) targeted the positive strand of the 171bp-duplicated integration sequence, with forward primer 5′-TTAGTGTCTTCCGGGTTCGC-3′, reverse primer 5′-GCGTTTCCTCACCACGTGA-3′, and probe 5′-/56-FAM/CAAGATGGC/ZEN/TGCCGTGACGTCACTAATACG/3IABkFQ/-3′. A reference RPP30 primer/probe set was used, with forward primer 5′-AGATTTGGACCTGCGAGCG-3′, reverse primer 5′-GAGCGGCTGTCTCCACAAGT-3′, and probe 5′- VIC/ TTCTGACCTGAAGGCTCTGCGCG −3′.

The ddPCR or RT-ddPCR was performed the kits ddPCR Supermix for Probes and One-Step RT-ddPCR Advanced Kit for Probes as recommended by manufacturer including 1 μl of nucleic acid in a final volume of 25 μl(Bio-Rad). For ddPCR reaction mix we included 1 μL of HindIII (New England Biolabs). Droplet generation was performed using a Bio-Rad automated droplet generator (Droplet Generation Oil for Probes). Amplification was carried out on a C1000 Thermal Cycler (Bio-Rad) with the following conditions: 95 °C for 10 min, 40 cycles of 94 °C for 30 s, 60 °C for 1 min, and a final 10-min hold at 98 °C. Droplets were analyzed on a QX200 or QX600 droplet reader (Bio-Rad) immediately following amplification. Data were processed with QX Manager Software (version 2.1), and quantification was calculated as copies/μL of specimen.

### Long-read sequencing of SKNO-1 cell lines

Frozen SKNO-1 cell pellets (5 x 10^6^ cells each) from the DSMZ and JCRB were submitted to the Translational Genomics Research Institute (TGen) for high-molecular-weight DNA extraction using the NEB Monarch HMW Extraction Kit for Cells & Blood. DNA quality was assessed using the Agilent Femto Pulse, followed by shearing using the Megaruptor 3 (Diagenode). Library preparation was performed using the SMRT-Bell Prep kit V3, and sequencing was carried out on the PacBio Revio system using SMRT-Cell technology, targeting an estimated 40X coverage for the human genome from each cell line.

Long reads were initially mapped to the de novo assembled SKNO-1 anellovirus isolate using minimap2 version 2.29-r1283 with the preset “-ax map-hifi” for high fidelity reads from PacBio. Supplementary and unmapped reads were filtered out with samtools version 1.21 (56). Reads were then aligned to the human genome GRCh38 analysis set (GCA_000001405.15, https://hgdownload.soe.ucsc.edu/goldenPath/hg38/bigZips/analysisSet/hg38.analysisSet.chroms.tar.gz).

Mapping of long reads to the human genome was performed using minimap2 with the preset-ax map-hifi, and the following parameters: -t 10 -L --no-supplementary −O 3,18 -E 2,1, adjusting gap penalties to improve indel resolution. Only primary alignments were retained for downstream analysis. Depth of coverage for mapping assemblies and specific genes was recorded using PanDepth version 2.26 (57).

### Motif Search and Statistical Analysis

Transcription factor binding motifs within the integrated viral genome were identified using FIMO v5.5.9 (2025) within the MEME suite (33, 58) and relative score analysis using the JASPAR database v11 2026 (34). We downloaded the position-weighted matrices for human transcription factors MAX, SPI-1/PU.1, ERG, FLI1, and ETV6 from JASPAR2026 and input them into FIMO v5.5.9 (https://meme-suite.org/meme/tools/fimo) along with the first 312 bp of the anellovirus integrant and used a threshold p-value < 10^-4^. JASPAR analysis was conducted using the same position-weighted matrices and 312 bp anellovirus sequence using the scan option (https://jaspar.elixir.no/) and a relative profile score threshold of >80%. Full results for each analysis are presented in Supplementary Tables 6 and 7.

### Phylogenetic analysis

We used reference anellovirus genomes provided by ICTV (https://ictv.global/vmr) for phylogenetic analysis. The ORF1 coding sequence of all representative *Alphatorquevirus*, *Betatorquevirus*, and *Gammatorquevirus* species reported by ICTV were downloaded from NCBI GenBank (accessed October 2, 2025). In addition, a set of 7 anellovirus genomes with over 90% identity by BLASTn to the assembled SKNO-1 anellovirus genome were also included. Genomes were trimmed to the annotated ORF1 coding sequence and aligned with MAFFT v7.526. ORF1 sequences were also translated based on the standard code using AliView v1.28 and aligned with MAFFT-DASH v7.526. Phylogenetic trees of the nucleotide and amino acid alignments of ORF1 were constructed with IQ-TREE v3.0.1 with 1,000 replicates of ultrafast bootstrap. Trees were visualized with Figtree.

### Protein structure prediction

We downloaded the Cryo-EM structures of Betatorquevirus LY1 anellovirus virus-like particle expressed in HEK293 from PDB (ID: 8V7X, resolution: 2.80Å, and 8CYG, resolution 3.98Å) (36). In addition, we downloaded the nucleotide sequence of Betatorquevirus LY1 ORF1 (NCBI GenBank: JX134044.1) and translated using the standard code. The Alphatorquevirus sp. isolate SKNO-1 ORF1 amino acid sequence was translated from the longest coding sequence of assembled genome. We used the amino acid sequences to predicted the structure using AlphaFold3 Server (accessed 2025-11-17) (59). The best prediction model (pTM = 0.84) provided by AlphaFold3 was used for comparative visualization in ChimeraX-1.8 (60).

## Data availability

NCBI SRA accession number analyzed in this study are available in Supplementary Tables S1-3. The sequenced Alphatorquevirus sp. isolate SKNO-1 genome was submitted to NCBI GenBank with accession number PQ220294.2. PacBio sequencing reads containing the anellovirus integrant were uploaded to NCBI SRA with accession numbers SRR36563182 and SRR36563183 (BioProject: PRJNA1392521).

## ACKNOWLEDGMENTS

The authors thank Yael L. Perez, Shah A.M. Bakhash, Kevin Kong, and Jeffrey Furlong for technical assistance. This research received no specific grant from any funding agency in the public, commercial, or not-for-profit sectors. ALG reports contract testing to UW from Abbott, Cepheid, Novavax, Pfizer, Janssen, Assembly Biosciences, Aicuris, Innovative Molecules, and Hologic, research support from Gilead, and personal fees from Arisan Therapeutics, outside of the described work.

## REFERENCES

1. Nishizawa T, Okamoto H, Konishi K, Yoshizawa H, Miyakawa Y, Mayumi M. 1997. A novel DNA virus (TTV) associated with elevated transaminase levels in posttransfusion hepatitis of unknown etiology. Biochem Biophys Res Commun 241:92–97.

2. Kumata R, Ito J, Takahashi K, Suzuki T, Sato K. 2020. A tissue level atlas of the healthy human virome. BMC Biol 18:55.

3. Sabbaghian M, Gheitasi H, Shekarchi AA, Tavakoli A, Poortahmasebi V. 2024. The mysterious anelloviruses: investigating its role in human diseases. BMC Microbiology 24:40.

4. Arze CA, Springer S, Dudas G, Patel S, Bhattacharyya A, Swaminathan H, Brugnara C, Delagrave S, Ong T, Kahvejian A, Echelard Y, Weinstein EG, Hajjar RJ, Andersen KG, Yozwiak NL. 2021. Global genome analysis reveals a vast and dynamic anellovirus landscape within the human virome. Cell Host & Microbe 29:1305–1315.e6.

5. Abbas AA, Young JC, Clarke EL, Diamond JM, Imai I, Haas AR, Cantu E, Lederer DJ, Meyer K, Milewski RK, Olthoff KM, Shaked A, Christie JD, Bushman FD, Collman RG. 2019. Bidirectional transfer of anelloviridae lineages between graft and host during lung transplantation. Am J Transplant 19:1086–1097.

6. Hamad Y, Charya A, Kong H, Jang M, Andargie T, Shah P, Mathew J, Orens J, Aryal S, Nathan S, Agbor-Enoh S. 2023. (1207) Anellovirus: A Novel Marker for Overimmunosuppression and Risk of Infection in Lung Transplant Recipients. The Journal of Heart and Lung Transplantation 42:S516.

7. Kelly E, Awan A, Sweeney C, Wildes D, De Gascun C, Hassan J, Riordan M. 2024. Torque Teno Virus Loads as a Marker of Immunosuppression in Pediatric Kidney Transplant Recipients. Pediatr Transplant 28:e14857.

8. Uyanik-Uenal K, Perkmann-Nagele N, Wittmann F, Ceran E, Aliabadi-Zuckermann A, Laufer G, Puchhammer-Stoeckl E, Zuckermann A. 2020. Torque Teno Virus DNA Load after Heart Transplantation and Its Association with the Strength of Immunosuppression: Preliminary Data of a Prospective Single Center Study. The Journal of Heart and Lung Transplantation 39:S128.

9. Engel B, Görzer I, Campos-Murguia A, Hartleben B, Puchhammer-Stöckl E, Jaeckel E, Taubert R. 2023. Association of torque teno virus viremia with liver fibrosis in the first year after liver transplantation. Front Immunol 14:1215868.

10. Boisvert N, Thurmond S, Elenberger C, Jeraldo P, Prince C, Sutherland N, Melo J, Tsheowang K, Dodier C, Nogalski M, Verma D, Parsons G, Cabral J. 2025. Anellovirus protein coded by ORF2/3 recruits host cell replication and homologous recombination machinery during replication. bioRxiv 10.1101/2025.05.28.656439.

11. Butkovic A, Kraberger S, Smeele Z, Martin DP, Schmidlin K, Fontenele RS, Shero MR, Beltran RS, Kirkham AL, Aleamotu’a M, Burns JM, Koonin EV, Varsani A, Krupovic M. 2023. Evolution of anelloviruses from a circovirus-like ancestor through gradual augmentation of the jelly-roll capsid protein. Virus Evol 9:vead035.

12. Kaczorowska J, Timmerman AL, Deijs M, Kinsella CM, Bakker M, van der Hoek L. 2023. Anellovirus evolution during long-term chronic infection. Virus Evolution 9:vead001.

13. Kraberger S, Serieys LEK, Richet C, Fountain-Jones NM, Baele G, Bishop JM, Nehring M, Ivan JS, Newkirk ES, Squires JR, Lund MC, Riley SPD, Wilmers CC, van Helden PD, Van Doorslaer K, Culver M, VandeWoude S, Martin DP, Varsani A. 2021. Complex evolutionary history of felid anelloviruses. Virology 562:176–189.

14. De Koch MD, Krupovic M, Fielding R, Smith K, Schiavone K, Hall KR, Reid VS, Boyea D, Smith EL, Schmidlin K, Fontenele RS, Koonin EV, Martin DP, Kraberger S, Varsani A. 2024. Novel lineage of anelloviruses with large genomes identified in dolphins. Journal of Virology 99:e01370–24.

15. Paietta EN, Kraberger S, Lund MC, Vargas KL, Custer JM, Ehmke E, Yoder AD, Varsani A. 2024. Diverse Circular DNA Viral Communities in Blood, Oral, and Fecal Samples of Captive Lemurs. Viruses 16:1099.

16. de Souza WM, Fumagalli MJ, de Araujo J, Sabino-Santos G, Maia FGM, Romeiro MF, Modha S, Nardi MS, Queiroz LH, Durigon EL, Nunes MRT, Murcia PR, Figueiredo LTM. 2018. Discovery of novel anelloviruses in small mammals expands the host range and diversity of the Anelloviridae. Virology 514:9–17.

17. Bigarré L, Beven V, de Boisséson C, Grasland B, Rose N, Biagini P, Jestin A. 2005. Pig anelloviruses are highly prevalent in swine herds in France. J Gen Virol 86:631–635.

18. Kakkola L, Bondén H, Hedman L, Kivi N, Moisala S, Julin J, Ylä-Liedenpohja J, Miettinen S, Kantola K, Hedman K, Söderlund-Venermo M. 2008. Expression of all six human Torque teno virus (TTV) proteins in bacteria and in insect cells, and analysis of their IgG responses. Virology 382:182–189.

19. Kamahora T, Hino S, Miyata H. 2000. Three spliced mRNAs of TT virus transcribed from a plasmid containing the entire genome in COS1 cells. J Virol 74:9980–9986.

20. Qiu J, Kakkola L, Cheng F, Ye C, Söderlund-Venermo M, Hedman K, Pintel DJ. 2005. Human Circovirus TT Virus Genotype 6 Expresses Six Proteins following Transfection of a Full-Length Clone. J Virol 79:6505–6510.

21. Kaczorowska J, van der Hoek L. 2020. Human anelloviruses: diverse, omnipresent and commensal members of the virome. FEMS Microbiol Rev 44:305–313.

22. Maggi F, Fornai C, Zaccaro L, Morrica A, Vatteroni ML, Isola P, Marchi S, Ricchiuti A, Pistello M, Bendinelli M. 2001. TT virus (TTV) loads associated with different peripheral blood cell types and evidence for TTV replication in activated mononuclear cells. Journal of Medical Virology 64:190–194.

23. Zhong S, Yeo W, Tang M, Liu C, Lin X, Ho WM, Hui P, Johnson PJ. 2002. Frequent detection of the replicative form of TT virus DNA in peripheral blood mononuclear cells and bone marrow cells in cancer patients. J Med Virol 66:428–434.

24. de Villiers E-M, Borkosky SS, Kimmel R, Gunst K, Fei J-W. 2011. The Diversity of Torque Teno Viruses: In Vitro Replication Leads to the Formation of Additional Replication-Competent Subviral Molecules. Journal of Virology 85:7284–7295.

25. Matozaki S, Nakagawa T, Kawaguchi R, Aozaki R, Tsutsumi M, Murayama T, Koizumi T, Nishimura R, Isobe T, Chihara K. 1995. Establishment of a myeloid leukaemic cell line (SKNO-1) from a patient with t(8;21) who acquired monosomy 17 during disease progression. Br J Haematol 89:805–811.

26. Greenblatt SM, Man N, Hamard P-J, Asai T, Karl D, Martinez C, Bilbao D, Stathias V, Jermakowicz AM, Duffort S, Tadi M, Blumenthal E, Newman S, Vu L, Xu Y, Liu F, Schurer SC, McCabe MT, Kruger RG, Xu M, Yang F-C, Tenen DG, Watts J, Vega F, Nimer SD. 2018. CARM1 Is Essential for Myeloid Leukemogenesis but Dispensable for Normal Hematopoiesis. Cancer Cell 33:1111–1127.e5.

27. Mas G, Man N, Nakata Y, Martinez-Caja C, Karl D, Beckedorff F, Tamiro F, Chen C, Duffort S, Itonaga H, Mookhtiar AK, Kunkalla K, Valencia AM, Collings CK, Kadoch C, Vega F, Kogan SC, Shiekhattar R, Morey L, Bilbao D, Nimer SD. 2023. The SWI/SNF chromatin-remodeling subunit DPF2 facilitates NRF2-dependent antiinflammatory and antioxidant gene expression. J Clin Invest 133:e158419.

28. Shah V, Giotopoulos G, Osaki H, Meyerhöfer M, Meduri E, Gallego-Crespo A, Behrendt MA, Saura-Pañella M, Tarkar A, Schubert B, Yun H, Horton SJ, Agrawal-Singh S, Haehnel PS, Basheer F, Lugo D, Eleftheriadou I, Barbash O, Dhar A, Kühn MWM, Guezguez B, Theobald M, Kindler T, Gallipoli P, Yeh P, Dawson MA, Prinjha RK, Huntly BJP, Sasca D. 2025. Acute resistance to BET inhibitors remodels compensatory transcriptional programs via p300 coactivation. Blood 145:748–764.

29. Harada T, Heshmati Y, Kalfon J, Perez MW, Xavier Ferrucio J, Ewers J, Hubbell Engler B, Kossenkov A, Ellegast JM, Yi JS, Bowker A, Zhu Q, Eagle K, Liu T, Kai Y, Dempster JM, Kugener G, Wickramasinghe J, Herbert ZT, Li CH, Vrabič Koren J, Weinstock DM, Paralkar VR, Nabet B, Lin CY, Dharia NV, Stegmaier K, Orkin SH, Pimkin M. 2022. A distinct core regulatory module enforces oncogene expression in KMT2A-rearranged leukemia. Genes Dev 36:368–389.

30. Paudel BB, Tan S-F, Fox TE, Ung J, Golla U, Shaw JJP, Dunton W, Lee I, Fares WA, Patel S, Sharma A, Viny AD, Barth BM, Tallman MS, Cabot M, Garrett-Bakelman FE, Levine RL, Kester M, Feith DJ, Claxton D, Janes KA, Loughran TP. 2024. Acute myeloid leukemia stratifies as 2 clinically relevant sphingolipidomic subtypes. Blood Adv 8:1137–1142.

31. Quentmeier H, Pommerenke C, Dirks WG, Eberth S, Koeppel M, MacLeod RAF, Nagel S, Steube K, Uphoff CC, Drexler HG. 2019. The LL-100 panel: 100 cell lines for blood cancer studies. Sci Rep 9:8218.

32. Kalantar KL, Carvalho T, de Bourcy CFA, Dimitrov B, Dingle G, Egger R, Han J, Holmes OB, Juan Y-F, King R, Kislyuk A, Lin MF, Mariano M, Morse T, Reynoso LV, Cruz DR, Sheu J, Tang J, Wang J, Zhang MA, Zhong E, Ahyong V, Lay S, Chea S, Bohl JA, Manning JE, Tato CM, DeRisi JL. 2020. IDseq—An open source cloud-based pipeline and analysis service for metagenomic pathogen detection and monitoring. GigaScience 9:giaa111.

33. Grant CE, Bailey TL, Noble WS. 2011. FIMO: scanning for occurrences of a given motif. Bioinformatics 27:1017–1018.

34. Rauluseviciute I, Riudavets-Puig R, Blanc-Mathieu R, Castro-Mondragon JA, Ferenc K, Kumar V, Lemma RB, Lucas J, Chèneby J, Baranasic D, Khan A, Fornes O, Gundersen S, Johansen M, Hovig E, Lenhard B, Sandelin A, Wasserman WW, Parcy F, Mathelier A. 2024. JASPAR 2024: 20th anniversary of the open-access database of transcription factor binding profiles. Nucleic Acids Res 52:D174–D182.

35. Varsani A, Kraberger S, Opriessnig T, Maggi F, Celer V, Okamoto H, Biagini P. 2023. Anelloviridae taxonomy update 2023. Arch Virol 168:277.

36. Liou S-H, Boggavarapu R, Cohen NR, Zhang Y, Sharma I, Zeheb L, Mukund Acharekar N, Rodgers HD, Islam S, Pitts J, Arze C, Swaminathan H, Yozwiak N, Ong T, Hajjar RJ, Chang Y, Swanson KA, Delagrave S. 2024. Structure of anellovirus-like particles reveal a mechanism for immune evasion. Nat Commun 15:7219.

37. Cordey S, Laubscher F, Hartley M-A, Junier T, Keitel K, Docquier M, Guex N, Iseli C, Vieille G, Le Mercier P, Gleizes A, Samaka J, Mlaganile T, Kagoro F, Masimba J, Said Z, Temba H, Elbanna GH, Tapparel C, Zanella M-C, Xenarios I, Fellay J, D’Acremont V, Kaiser L. 2021. Blood virosphere in febrile Tanzanian children. Emerg Microbes Infect 10:982–993.

38. Cabezas-Wallscheid N, Eichwald V, de Graaf J, Löwer M, Lehr H-A, Kreft A, Eshkind L, Hildebrandt A, Abassi Y, Heck R, Dehof AK, Ohngemach S, Sprengel R, Wörtge S, Schmitt S, Lotz J, Meyer C, Kindler T, Zhang D-E, Kaina B, Castle JC, Trumpp A, Sahin U, Bockamp E. 2013. Instruction of haematopoietic lineage choices, evolution of transcriptional landscapes and cancer stem cell hierarchies derived from an AML1-ETO mouse model. EMBO Mol Med 5:1804–1820.

39. Al-Harbi S, Aljurf M, Mohty M, Almohareb F, Ahmed SOA. 2020. An update on the molecular pathogenesis and potential therapeutic targeting of AML with t(8;21)(q22;q22.1);RUNX1-RUNX1T1. Blood Adv 4:229–238.

40. Astolfi A, Masetti R, Indio V, Bertuccio SN, Messelodi D, Rampelli S, Leardini D, Carella M, Serravalle S, Libri V, Bandini J, Volinia S, Candela M, Pession A. 2021. Torque teno mini virus as a cause of childhood acute promyelocytic leukemia lacking PML/RARA fusion. Blood 138:1773–1777.

41. Sala-Torra O, Beppu LW, Abukar FA, Radich JP, Yeung CCS. 2022. TTMV-RARA fusion as a recurrent cause of AML with APL characteristics. Blood Adv 6:3590–3592.

42. Sun S, Liu Y, Xu Q-Y, Zhou X, Wen L, Chen L, Chen J, Wang H, Chen X, Lou J, Zheng H, Xie Y, Wang L, Ma F, Yang L, Wang X, Gao W, Wu Y, Zhang L, Shen S, Chen S, Wang K, Liu H, Huang J, Zhu H-H. 2024. Comprehensive Clinical and Molecular Characterization Reveals Rara As the Most Frequent Torque Teno Mini Virus Insertion Site and Defines a Distinct Ttmv::Rara Driven APL-like Leukemia Subtype. Blood 144:1547.

43. Kosulin K, Kernbichler S, Pichler H, Lawitschka A, Geyeregger R, Witt V, Lion T. 2018. Post-transplant Replication of Torque Teno Virus in Granulocytes. Front Microbiol 9:2956.

44. Albert E, Giménez E, Hernani R, Piñana JL, Solano C, Navarro D. 2024. Torque Teno Virus DNA Load in Blood as an Immune Status Biomarker in Adult Hematological Patients: The State of the Art and Future Prospects. Viruses 16:459.

45. Wang X-C, Wang H, Tan S-D, Yang S-X, Shi X-F, Zhang W. 2020. Viral metagenomics reveals diverse anelloviruses in bone marrow specimens from hematologic patients. Journal of Clinical Virology 132:104643.

46. Toppinen M, Sajantila A, Pratas D, Hedman K, Perdomo MF. 2021. The Human Bone Marrow Is Host to the DNAs of Several Viruses. Front Cell Infect Microbiol 11.

47. Suzuki T, Suzuki R, Li J, Hijikata M, Matsuda M, Li T-C, Matsuura Y, Mishiro S, Miyamura T. 2004. Identification of Basal Promoter and Enhancer Elements in an Untranslated Region of the TT Virus Genome. Journal of Virology 78:10820–10824.

48. Kamada K, Kamahora T, Kabat P, Hino S. 2004. Transcriptional regulation of TT virus: promoter and enhancer regions in the 1.2-kb noncoding region. Virology 321:341–348.

49. Li W, Wang S, Jin Y, Mu X, Guo Z, Qiao S, Jiang S, Liu Q, Cui X. 2024. The role of the hepatitis B virus genome and its integration in the hepatocellular carcinoma. Front Microbiol 15.

50. Gilbert-Girard S, Gravel A, Artusi S, Richter SN, Wallaschek N, Kaufer BB, Flamand L. 2017. Stabilization of Telomere G-Quadruplexes Interferes with Human Herpesvirus 6A Chromosomal Integration. J Virol 91:e00402–17.

51. Chatterjee S, Starrett GJ. 2024. Microhomology-mediated repair machinery and its relationship with HPV-mediated oncogenesis. Journal of Medical Virology 96:e29674.

52. Wahls WP, Wallace LJ, Moore PD. 1990. The Z-DNA motif d(TG)30 promotes reception of information during gene conversion events while stimulating homologous recombination in human cells in culture. Mol Cell Biol 10:785–793.

53. Zheng B, Gold SR, Iwanaszko M, Howard BC, Wang L, Shilatifard A. 2023. Distinct layers of BRD4-PTEFb reveal bromodomain-independent function in transcriptional regulation. Mol Cell 83:2896–2910.e4.

54. Wood DE, Lu J, Langmead B. 2019. Improved metagenomic analysis with Kraken 2. Genome Biol 20:257.

55. Bankevich A, Nurk S, Antipov D, Gurevich AA, Dvorkin M, Kulikov AS, Lesin VM, Nikolenko SI, Pham S, Prjibelski AD, Pyshkin AV, Sirotkin AV, Vyahhi N, Tesler G, Alekseyev MA, Pevzner PA. 2012. SPAdes: a new genome assembly algorithm and its applications to single-cell sequencing. J Comput Biol 19:455–477.

56. Li H. 2018. Minimap2: pairwise alignment for nucleotide sequences. Bioinformatics 34:3094–3100.

57. Yu H, Shi C, He W, Li F, Ouyang B. 2024. PanDepth, an ultrafast and efficient genomic tool for coverage calculation. Briefings in Bioinformatics 25:bbae197.

58. Bailey TL, Johnson J, Grant CE, Noble WS. 2015. The MEME Suite. Nucleic Acids Res 43:W39–W49.

59. Abramson J, Adler J, Dunger J, Evans R, Green T, Pritzel A, Ronneberger O, Willmore L, Ballard AJ, Bambrick J, Bodenstein SW, Evans DA, Hung C-C, O’Neill M, Reiman D, Tunyasuvunakool K, Wu Z, Žemgulytė A, Arvaniti E, Beattie C, Bertolli O, Bridgland A, Cherepanov A, Congreve M, Cowen-Rivers AI, Cowie A, Figurnov M, Fuchs FB, Gladman H, Jain R, Khan YA, Low CMR, Perlin K, Potapenko A, Savy P, Singh S, Stecula A, Thillaisundaram A, Tong C, Yakneen S, Zhong ED, Zielinski M, Žídek A, Bapst V, Kohli P, Jaderberg M, Hassabis D, Jumper JM. 2024. Accurate structure prediction of biomolecular interactions with AlphaFold 3. Nature 630:493–500.

60. Meng EC, Goddard TD, Pettersen EF, Couch GS, Pearson ZJ, Morris JH, Ferrin TE. 2023. UCSF ChimeraX: Tools for structure building and analysis. Protein Science 32:e4792.

